# Sea lettuce systematics: lumping or splitting?

**DOI:** 10.1101/413450

**Authors:** Manuela Bernardes Batista, Regina L. Cunha, Rita Castilho, Paulo Antunes Horta

**Author notes:** These authors contributed equally to the work (Manuela Bernardes Batista, Regina L. Cunha).

## Abstract

Phylogenetic relationships within sea lettuce species belonging to the genus *Ulva* is a daunting challenge given the scarcity of diagnostic morphological features and the pervasive phenotypic plasticity. With more than 100 species described on a morphological basis, an accurate evaluation of its diversity is still missing. Here we analysed 277 chloroplast-encoded gene sequences (43 from this study), representing 35 nominal species of *Ulva* from the Pacific, Indian Ocean, and Atlantic (with a particular emphasis on the Brazilian coast) in an attempt to solve the complex phylogenetic relationships within this widespread genus. Maximum likelihood, Bayesian analyses and species delimitation tests support the existence of 22 evolutionary significant units (ESUs), lumping the currently recognized number of species. All individuals sampled throughout an extensive area of the Brazilian coast were included within two distinct ESUs. Most of the clades retrieved in the phylogenetic analyses do not correspond to a single nominal species. Geographic range evolution indicated that the ancestor of *Ulva* had a distribution restricted to the temperate North Pacific. The dating analysis estimated its origin during the Upper Cretaceous at 75.8 million years (myr) ago but most of the cladogenetic events within the genus occurred in the last ten myr. Pervasive human-mediated gene flow through ballast water and widespread morphologic plasticity are the most likely explanations for the difficulty in establishing a reliable phylogenetic framework for this conspicuous, widespred and many times abundant green algae morphotype.

## Introduction

The genus *Ulva* Linnaeus (Ulvophyceae, Ulvales) commonly known as sea lettuce is gaining global relevance in coastal ecosystem management owing to their increased role in generating green tides [1]. With the increasing need of targeting species for restoration protocols, the precise identification of fast growing autochthonous species is critical for this process. *Ulva* is a ubiquitous macroalgal genus showing species inhabiting a remarkable variation of aquatic environments ranging from shallow rocky shores to the subtidal, up to 100 meters deep [2]. Due to their high salinity tolerance, some *Ulva* species may grow under marine, estuarine or brackish conditions. They can either be found in areas strongly impacted by anthropogenic activities or in pristine habitats [3,4]. Sea lettuce species are among the most abundant organisms attached to ship hulls and ballast water [5]. Some species exhibit invasive traits (e.g., high growth rates or efficient reproductive alternatives) that promote dispersal abilities [6].

The systematics of this group of marine macroalgae remains challenging given their morphological simplicity [7], widespread plasticity, intraspecific variation, and interspecific overlap [8]. With more than 100 nominal species described on a morphological basis [9], several are considered homotypic or heterotypic synonyms [10], and a precise evaluation of current biodiversity within the genus *Ulva* is not yet available. This morphologic framework with many potential cryptic species also precludes robust biogeographic interpretations and fails to recognize connectivity and dispersal processes, essential for ecosystem management [11].

Molecular tools have been an important ally not only for tracking potentially harmful “green tide” species but also represent a more accurate procedure to evaluate the taxonomy and cryptic diversity within *Ulva* [12]. Thus far, molecular studies of the genus focused on more restricted geographic areas (temperate and boreal waters of Europe [9]; north-western American coast [13,14,15]; New Zealand [16], southern Australia [17]; Hawaii [18,19]; western coast of India [12], and Japan [20]). Although the study of Hayden and colleagues [13] includes 21 *Ulva* species, it is focused on the North American coast and a worldwide phylogeny is still missing. Further, the genetic assessment of *Ulva* species from the southwestern Atlantic, particularly of the Brazilian coast is completely absent. This region may hold fundamental information to complete the systematic relationships within *Ulva* and complete the biogeographical puzzle of the sea lettuce representatives, considering the global processes of dispersion, invasion, and connectivity occurring in this area [21,22,23]. From a global perspective, seaweed phylogeographic studies promote a better understanding on how environmental processes along an ecological and evolutionary time scale influence complex patterns of inter- and intraspecific genetic diversity, fostering discussions regarding lumping or splitting systematics[24,25].

In this study, we analysed all available sequences (277 in total, 43 from this study) of the large subunit of the chloroplast-encoded RUBISCO gene (*rbcL*), representing 35 nominal species of *Ulva* with distromatic blade-form from the Pacific, Indian Ocean, and Atlantic (with a particular emphasis on the Brazilian coast) in an attempt to better understand the complex phylogenetic relationships within this widespread genus. We performed three different species delimitation tests to better evaluate lineage boundaries, and estimated major lineage splitting events within *Ulva* to infer its evolutionary history using a Bayesian approach.

## Results

### Species delimitation tests

Species delimitation tests grounded on molecular data arrived at different results. GMYC indicated the existence of 22 evolutionary significant units (ESUs) while mPTP and ABGD indicated 13 and 10 ESUs, respectively (Fig. 2). The northern hemisphere temperate regions hosted the higher number of ESUs (endemic and non-endemic) and unique haplotypes. Overall, species recovered by species delimitation tests coincided with well-supported clades in both ML and BI analyses. *Ulva limnetica* was recovered as an ESU by the three methods while *Ulva arasakii* was identified as a valid ESU by GMYC and mPTP, only. The GYMC method is based on an ultrametric tree, here produced by BEAST, which does not accommodate unresolved nodes, therefore, it is expectable that the recovered ESUs do not correspond to the ML clades.

**Figure 1.**
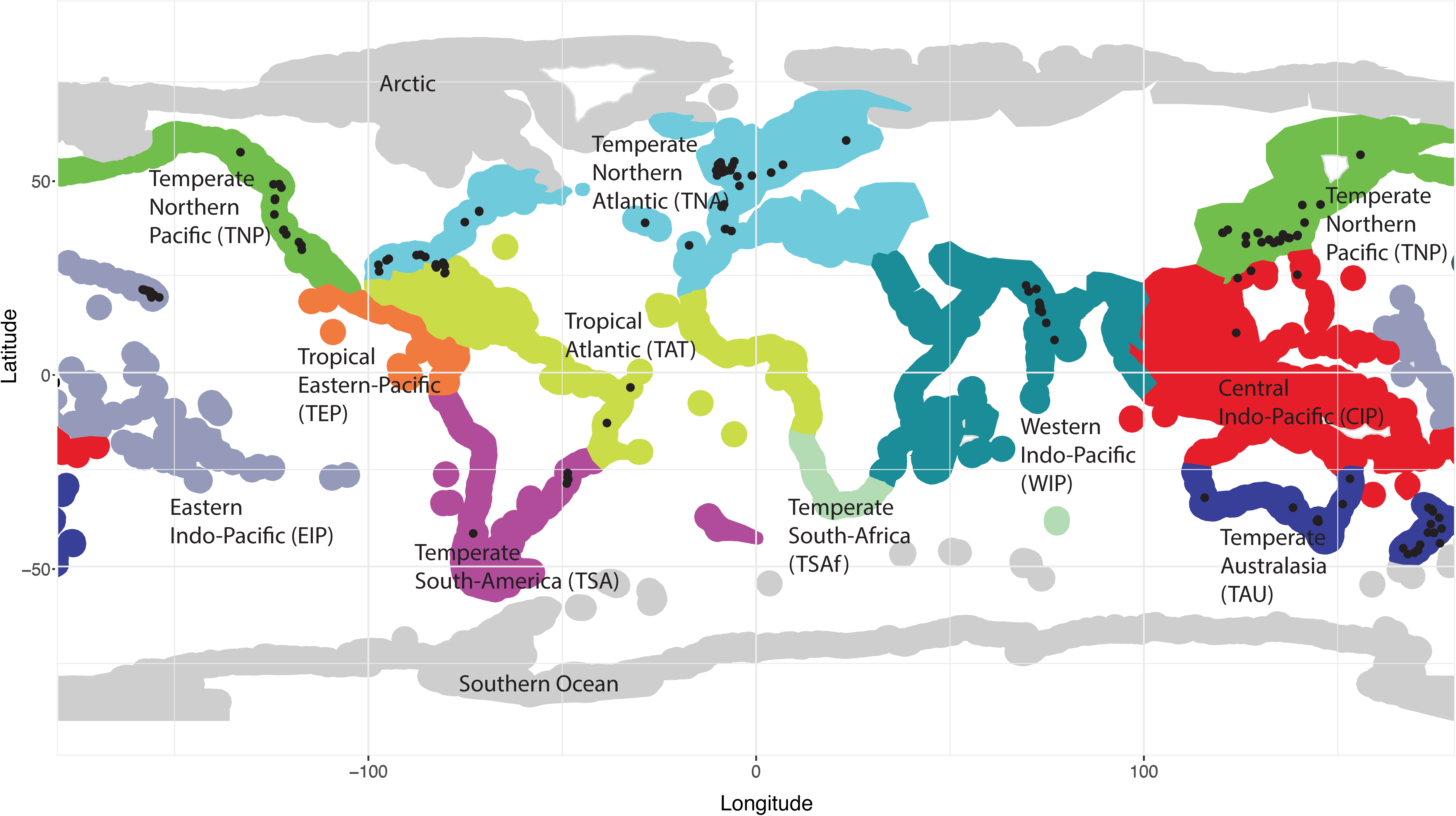
Biogeographic realms according to [65].

**Figure 2.**
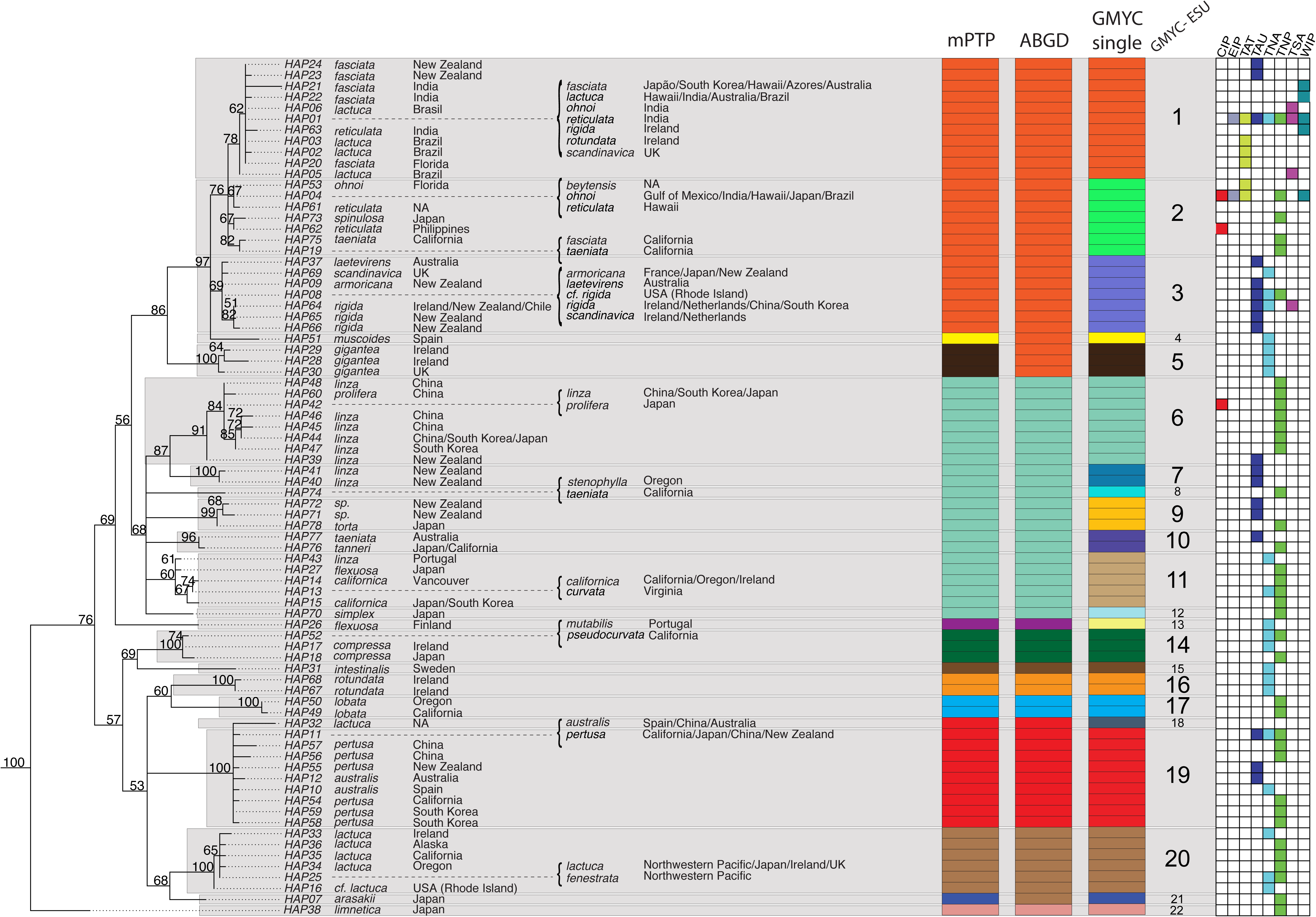
Phylogenetic relationships of *Ulva* based on maximum analysis of a portion of the large subunit of the chloroplast-encoded RUBISCO gene (*rbcL*) produced by PhyML. Numbers at the nodes represent bootstrap proportions. Assignments of each *Ulva* sample to delineated entities from mPTP, ABGD and GMYC species delimitation tests are shown by coloured rectangles. The estimated Evolutionary Significant Units (ESUs) by GMYC are depicted in grey rectangles. Biogeographic realms according to [65] assigned to each sample are shown by colored squares (CIP, Central Indo-Pacific; EIP, Eastern Indo-Pacific; TAT, Tropical Atlantic; TAU, Temperate Australasia; TNA, Temperate North Atlantic; TNP, Temperate North Pacific; TSA, Temperate South America; WIP, Western Indo-Pacific.

*Ulva gigantea* was recovered as a valid ESU by mPTP and GMYC. Samples from the Brazilian coast morphologically identified as *U. lactuca* and *U. ohnoi* were assigned (in the GMYC analysis) to the ESU1 dominated by *U. fasciata* or to ESU 2, respectively.

### Phylogenetic analyses

ML analysis of the *rbcL* data set (277 individuals, 1141 bp) yielded the topology (-ln *L* = 3659.98) depicted in Fig. 2. BI analysis retrieved a similar tree (-ln *L* = 3883.26) with ESS (Effective Sample Sizes) much larger than 200, indicating convergence of the runs. Convergence diagnostic in Bayesian analyses indicates that as potential scaler factors (PSRF) approaches 1.0, runs converge [26]. PSRF of our Bayesian analysis were all 1.00 indicating run convergence.

The vast majority of the clades recovered by the ML analysis do not correspond to the currently recognized nominal species within the genus but are congruent with results from GMYC (Fig. 2). The first main clade (Bootstrap Proportions, BP = 78%), contains a group of 7 morphologically-defined taxa (*Ulva fasciata / reticulata / lactuca / rotundata / scandinavica / rigida / ohnoi*) comprehending most of the *Ulva*’s distributional range in the northern and southern hemispheres, as well as in tropical and temperate environments. This clade was identified as ESU 1 by GMYC. The second main clade (BP = 69%), corresponds to the nominal species *Ulva scandinavica / laetevirens / armoricana / rigida* and was recognized as ESU 3. The next clade (BP = 100%), corresponds to *U. gigantea* (ESU 5). The following main clade (BP = 91%) comprehends most of the specimens assigned to *U. linza* (ESU 6), except the two from New Zealand that group in a different clade (BP = 100%), and were identified as a different ESU (ESU 7). Another clade (BP = 99%) includes three specimens, one assigned to *U. torta* from Japan and two *Ulva* sp. from New Zealand, classified as ESU 9 by GMYC. The clade including a specimen assigned to *U. taeniata* and another to *U. tanneri* received high statistical support (BP = 96%) and was identified as ESU 10. The ESU 11 identified by GMYC corresponds to a clade including *U. linza / flexuosa/ californica*/ *curvata*/. The next clade (BP = 100%) comprehends *U. compressa/ pseudocurvata/ mutabilis* and was classified as ESU 14. *Ulva rotundata* and *U. lobata* are recovered as sister clades and identified as ESUs 16 and 17, respectively. The clade including most of specimens assigned to *U. pertusa* and *U. australis* and one specimen identified as *U. lactuca* (PB = 100%) was classified as ESU 19 by GMYC. The last main clade (PB = 100%), included most of the specimens morphologically-identified as *U. lactuca*.

*Ulva limnetica* was recovered as the most basal lineage within the genus, and as a valid taxon by the three species delimitation methods. The 47 specimens from Brazil, previously identified as *U. lactuca* and *U. ohnoi*, were represented by six different haplotypes that were all included within clades 1 and 2.

### Date estimates

We estimated the age of the most recent common ancestor of the genus (MRCA) *Ulva* during the Upper Cretaceous at 75.79 [59.6-93.8] myr. The estimated ages (in million years) for the most recent common ancestor of each ESU determined by GMYC are the following: ESU 1 - 6.91 [3.59-10.9]; ESU 2 - 8.97 [4.5-13.9]; ESU 3 - 7.23 [3.0-13.0]; ESU 4 - 20.11 [13.3-28.1]; ESU 5 - 8.14 [3.1-14.6]; ESU 6 - 9.61 [4.9-15.4]; ESU 7 - 2.01 [0.1-5.9]; ESU 8 - 36.07 [26.2-48.0]; ESU 9 - 3.75 [0.9-8.1]; ESU 10 - 2.02 [0.1-5.7]; ESU 11- 9.52 [4.3-16.0]; ESU 12- 17.54 [9.4-26.8]; ESU 13 - 58.41 [44.0-75.2]; ESU 14 - 4.89 [1.2-10.6]; ESU 15 - 44.63 [29.2-61.4]; ESU 16 - 2.09 [0.05-6.1]; ESU 17 - 1.83 [0.05-5.2]; ESU 18 - 7.41 [3.8-12.1]; ESU 19 - 10.97 [5.7-17.7]; ESU 20 5.96 [2.4-10.9]; ESU 21 - 26.66 [15.9-38.9], and ESU 22 - 75.79 [59.6-93.8]. The remaining estimates for the siphonous green algae included in this analysis are congruent with a previous study [27] for the deeper nodes but are more recent for the tips (Fig. 3).

**Figure 3.**
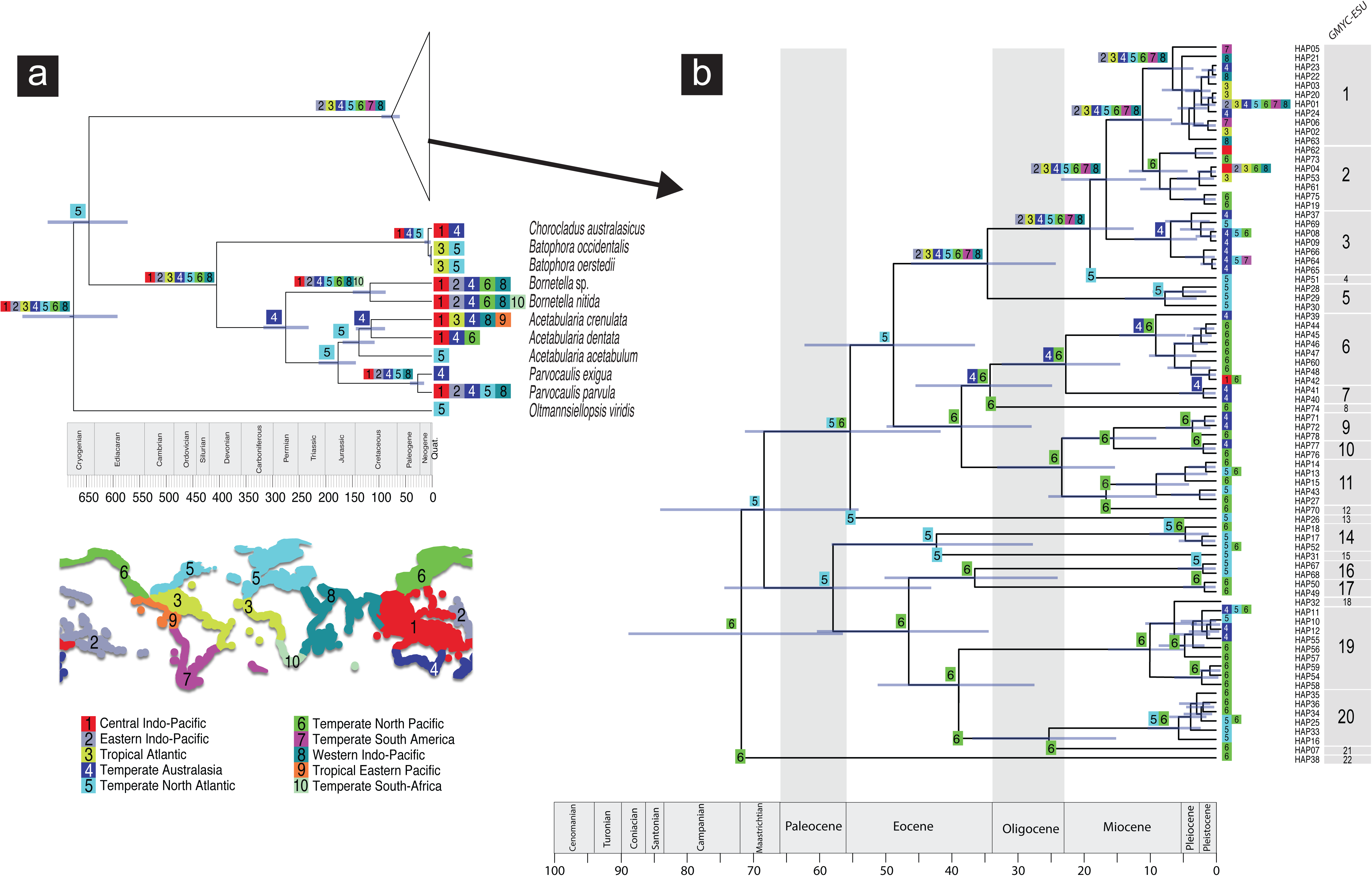
**a.** BEAST maximum clade credibility chronogram showing main cladogenetic events within the genus *Ulva* (in the cartoon) and several species of the order Dasycladales. Numbers at the coloured squares indicate present-day distribution of species at the tips, and ancestral ranges on the nodes. The corresponding 95% highest posterior density intervals (blue bars) are depicted; **b.** Detail of the cartoon showing main cladogenetic events within the genus *Ulva* estimated by BEAST with ancestral estimation inferred with BioGeoBEARS. Numbers at the coloured squares indicate present-day distribution of *Ulva* species at the tips, and ancestral ranges on the nodes. Map of the biogeographic realms according to [65] is also shown.

### Biogeographic analyses

The dispersal-extinction-cladogenesis (DEC) + “J” (jump dispersal or founder event speciation) was selected as the best-fitting biogeographic model for *Ulva* (-ln *L* = 274.4; Table 2). However, given the recent controversy involving the “J” parameter, we selected the second-best model DEC (-ln *L* = 286.8; Table 2). “DEC+J model is a poor model of founder-event speciation, and statistical comparisons of its likelihood with DEC are inappropriate” [28]. Further, the parameter “J” artificially inflates the contribution of cladogenetic events to the likelihood, and prevents of non-jump events at ancestral nodes [28].

The stem lineage of *Ulva* shows an ancestral area restricted to the temperate North Pacific (Fig. 3). Some of the estimated ESUs (GMYC) are confined to temperate Australasia (ESUs 3 and 7), temperate North Pacific (ESUs 2, 8, 9, 10, 11, 12, 17, 19 and 21) or temperate North Atlantic (ESUs 4, 5, 15, and 16), while ESU 1 showed the widest ancestral range covering the northern temperate regions of the Pacific and Atlantic, eastern and western Indo-Pacific and the tropical areas of the Atlantic and eastern Pacific (Fig. 3).

## Discussion

Previous studies focusing on relatively restricted geographic areas and a lower number of species have recurrently reported inconsistent intra-generic phylogenetic relationships within *Ulva sea lettuce representatives* [2,20]. Here, we revisited the systematics of the taxa with this morphotype analysing *rbcL* sequence data from 35 ‘nominal’ species, defined as “taxa named for our convenience” [29], representing almost its entire distribution.

Species delimitation tests indicated a variable number of ESUs depending on the method (ABGD: 10; mPTP: 13; GMYC: 22). Most of the species described on a morphological basis were not validated by mPTP and ABGD tests. For instance, ESU 1 identified both GMYC contains, among others, specimens assigned to *U. rigida, U. armoricana* and *U. scandinavica* (Fig. 2), a group of nominal species already identified as conspecific in a previous study based on *rbcL* sequence data and morphological information [9]. This study also concluded that the use of morphological characters in *Ulva* is“largely inconclusive and of limited value for the circumscription of species” [9]. Results presented here reinforce that phenotypic plasticity and overlapping diagnostic characters produce a complex scenario in which specimens with different morphoanatomical and structural features are included within the same ESU (Fig. 2).

ML and BI analyses were mostly consistent with previous studies using a fraction of our taxon sampling, but the inclusion of more species further complicated phylogenetic relationships within the genus. *Ulva gigantea* is morphologically identical to *U. pseudocurvata* but ML and BI analyses clearly separate both lineages (Fig. 2). On the contrary, and regardless their morphologic distinctiveness, molecular analyses indicated that specimens identified as *U. compressa* and *U. pseudocurvata* should be included within the same species (Fig. 2), as reported in previous studies [9,13,30,31]. Moreover, *U. mutabilis* shares the same haplotype as *U. pseudocurvata* (Fig. 2), and thus should be considered the same species. Our analyses showed that specimens identified as *U. californica* and *U. curvata* are conspecific, although considered a valid species in a previous study [13].

The combination of species delimitation tests and molecular phylogenetic analyses suggests the 22 ESUs indicated by GMYC, as a reasonable estimation of folios *Ulva* diversity. Molecular data and crossing-test experiments identified *U. fasciata* and *U. ohnoi* [5], as well as *U. flexuosa* and *U. californica* [32] as separate species. These findings corroborate results from the GMYC that was the only species delimitation method splitting *U. fasciata* and *U. ohnoi* into different ESUs (ESUs 1 and 2, respectively).

The inclusion of other molecular markers (e.g., nuclear ribosomal internal transcribed spacers, ITS or the elongation factor *tufA*) does not seem to add any relevant phylogenetic information to solve the complex relationships within *Ulva*. For instance, studies based on ITS showed that the nominal species *U. fasciata/U. armoricana/U. rigida/U. scandinavica/U. taeniata* [14,18] and *U. linza/U. prolifera/U. californica* [17] are found interspersed in the recovered clades as shown in phylogenies based on *rbcl* sequence data (present study; [9]). Another phylogenetic analysis of 17 *Ulva* species based on the elongation factor *tufa* retrieved similar phylogenetic relationships within the genus [33].

Temperate seas host most of the diversity of this group and include ESUs that are only present in one biogeographic region (e.g., ESUs 4, 5 and 16 from the North Atlantic; Fig. 3). This general pattern is supported by studies that characterise intermediate latitudes as more diverse regarding seaweeds [34]. However, there is some evidence of Chlorophyte high species richness also in the tropics, particularly in the Indo-Pacific region [35].

Ancestral range estimation showed that the MRCA of *Ulva* was distributed along the temperate North Pacific (Fig. 3). Most of the ESUs show ancestral distributions restricted to northern temperate regions of the Pacific or Atlantic, with two exceptions: (1) ESU 1 that is widely distributed in the northern temperate waters of the Pacific and Atlantic, temperate South America and Australasia, eastern and western Indo-Pacific, and (2) ESU 6 occurring both in the southern (temperate Australasia) and northern (temperate north Pacific) hemispheres. These findings suggest that *Ulva* ancestral lineage most likely adapted to distinct biogeographic regions and environmental conditions, which may have promoted diversification within the genus.

The crown group age of *Ulva* was estimated at 75.2 myr, but most of the cladogenetic events within the genus took place between 10 and two myr ago (Fig. 3), most likely experiencing the changes that took place in the northern hemisphere. The centre of origin (speciation) located in the North Pacific that comprehends most of the cold-temperate faunas was formed *c.* 40 myr ago, but only after the opening of the Bering Strait about 12 myr ago started what is called the Great Trans-Arctic Biotic Interchange [36]. The continuous biotic flow between the North Pacific and the Arctic-North Atlantic was fully established at about 3.5 myr and dramatically changed the ecosystems of the latter region [36]. When a major glaciation started between 2.9 and 2.4 myr ago, the cold-temperate biotas moved southward in both oceans (Pacific and Atlantic). The Great Trans-Arctic Biotic Interchange may explain the close phylogenetic affinity we found between ESUs belonging to the temperate-north Pacific and Atlantic biogeographic regions.

Current geographic distribution of the 22 identified ESUs reflects the existence of physiological barriers that may represent the main constraint for their distribution. However, there are ‘nominal species’ belonging to the same ESU that are from distinct biogeographic regions and environmental conditions, sharing the same haplotype (e.g., HAP01: *U. scandinavica* from the UK and *U. lactuca* from Australia), which suggests that temperature is not the only driver of *Ulva* distribution (Fig. 2). Natural dispersal in *Ulva* is mostly driven by the oceanic drifting of propagules or mature thalli detached from the substrate, or using floating substrates [37,38]. Therefore, the existence of individuals belonging to the same ESU and occurring in disjunct biogeographic regions (e.g., ESU 1; Fig. 3), suggests that anthropogenic transport may be playing an important role in their distribution.

The conflicting intra-generic phylogenetic relationships and the wide non-overlapping geographic distributions of some of the ESUs almost certainly result from the extensive human-mediated gene flow that can potentially confound species boundaries in more recently derived species, and an ancient age of the stem lineage of *Ulva* that may hinder phylogenetic information due to homoplasy. There is an increasing body of evidence showing that ballast water is an important factor on macroalgal transport [39]. Our reconstructed phylogenetic patterns with clades without any geographic correspondence and grouping samples from distant locations is consistent with a scenario of human-mediated dispersal. Clade 3, for instance, includes nine nominal species and groups samples from New Zealand, Portugal, Vancouver, Finland or Japan, which represent the world’s top trade routes for container shipping (https://geopoliticalfutures.com/top-container-ship-trade-routes/). Because *Ulva* is recognized as a suitable biological model to address topics such as tissue morphology, complex multicellularity and developmental biology [1] a well-established lineage phylogeny is pivotal to a sound foundation of those studies.

## Materials and Methods

### Sampling

The study area included an extensive area of the coast of Brazil (states of Santa Catarina, Parana, Bahia, and Fernando Noronha archipelago). We followed [40] for morphological identification. Species with tubular morphology were not sampled. Twenty individuals were randomly collected from the tidal region of each sampled locality, except in the Fernando de Noronha archipelago, Brazil where only two specimens were found. Samples were placed in individual plastic bags and transported to the laboratory in a refrigerated container. For the molecular analysis, a 2 × 2 cm fragment of the largest leave of each individual was removed and inserted in small tea bag to avoid crumbling and subsequently sealed in zip-lock plastic bag, filled by dry silica gel. All vouchers of the collected material are currently retained in the Laboratory of Phycology of the Federal University of Santa Catarina and will be deposited in the FLOR Herbarium. The list of specimens, sample location and GeneBank accession numbers are shown in the supplementary material_S1.

### DNA extraction, PCR amplification and Sequencing

Species identifications were based on both DNA sequence data and morphological information. About 10 mg of dried *Ulva* tissues were used for DNA extraction from the samples obtained in the Brazilian coast. Samples were placed in a 1.5 ml Eppendorf tube, frozen with liquid nitrogen and grounded for 2 minutes with a plastic pestle. DNA was extracted using a commercial kit [NucleoSpin^®^ Plant II, Macherey-Nagel, Düren, Germany] following the manufacturer’s instructions. The chloroplast-encoded *rbcL* gene was amplified in 47 specimens using the PCR profile described in [9] and the published primers pair RH1 and 1385r [41] that amplify the large subunit of the chloroplast-encoded RUBISCO gene (*rbcL*). After amplification, PCR products were purified using the kit PEG 8000 (Polyethyleneglycol 8000) [42].

The purified PCR products were subsequently sequenced by the chain termination method (Sanger et al., 1977), with the Big Dye Terminator v 3.1 kit (Thermo Scientific, Carlsbad, CA, USA) following the protocol specified by the supplier. Resulting cycle sequence reaction were purified with ETOH/EDTA precipitation and were either sequenced at the Universidade Federal de Santa Catarina, Laboratório de Fisiologia do Desenvolvimento e Genética Vegetal on a 3500 XL - Applied Biosystems (Thermo Scientific) or at the ABI 3730 - Applied Biosystems. Resulting chromatograms were assembled using the software Sequencing Analysis v6.0 (Thermo Scientific).

### Species delimitation tests

We used three different approaches to delimit evolutionary significant units (ESUs) within *Ulva*: (1) the Automatic Barcode Gap Discovery (ABGD) method [43]; (2) the single-rate Poisson Tree Processes method (PTP, [44]), and (3) the general mixed Yule-coalescent (GMYC) model. ABGD detects a gap in the distribution of pairwise distances and uses this information to partition the sequences into groups of hypothetical species. This analysis was performed through the web server of ABGD

http://wwwabi.snv.jussieu.fr/public/abgd/abgdweb.html), using three distance options (Jukes-Cantor, Kimura 2-parameter and Simple) and the remaining parameters were used as default. PTP considers that every species evolved at the same rate, using the within-species number of substitutions that are modelled with a single exponential distribution and the multi-rate Poisson Tree Processes (mPTP, [45]), which takes into account the different rates of branching events within each delimited species, assuming that the intraspecific coalescence rate may vary significantly even among sister species. GMYC uses a “threshold time before which all nodes reflect diversification events and after which all nodes reflect coalescent events” [46]. We used the single-threshold GMYC model as implemented in SPLITS, code written by T. Ezard, T. Fujisawa and T. Barraclough in R v.3.5 [47]. The required ultrametric tree was obtained with BEAST2 v2.5 [48] as described above.

All analyses were run on CCMAR computational cluster facility (http://gyra.ualg.pt)a and on R2C2 computational cluster facility (http://rcastilho.pt/R2C2/R2C2_cluster.html) both located at the IT Services of the University of Algarve.

### Phylogenetic Analyses

All DNA sequences were aligned using MAFFT v.7.245 (Multiple alignment using Fast Fourier Transform) [49] using the --auto option that automatically selects the appropriate strategy according to data size. The Akaike information criterion [50] as implemented in MODELTEST function of phangorn R-package [51] selected the GTR+I+Γ (I=0.6927; Γ=0.8877) as the model that fits better the data. These setting were used in the maximum likelihood (ML) analysis that was performed with PhyML v3.0 [52]. The robustness of the inferred trees was tested by nonparametric bootstrapping (BP) using 1000 pseudoreplicates. Bayesian inferences (BI) were performed with MRBAYES v.3.1.2 [53] by Metropolis coupled Markov chain Monte Carlo (MCMCMC) sampling for 20×10^6^ generations (four simultaneous MC chains; sample frequency 1000). Length of burn-in was determined by visual examination of traces in TRACER v.1.6 [54]. According to prior information [9], *Umbraulva olivascens* was selected as outgroup in both analyses.

### Dating Analysis

BEAST v.1.8.4 [55] was used to date the origin of the genus *Ulva*. We included several species of siphonous green algae from the order Dasycladales that have paleontological record to allow the calibration of the phylogeny. First calibration refers to the earliest occurrence of the fossil *Uncatoella verticillata* from the Lower Devonian [419.2-397.5] myr [56]. This fossil has the thallus and reproductive structures comparable to those of extant Dasycladaceae [57] here represented by the stem lineage of the clade *Batophora occidentalis, B. oerstedii* and *Chlorocladus australasicus*. This calibration was modeled with a normal distribution (mean=408.35, stdev=10.85). The second calibration refers to the fossil *Acicularia heberti* from the Danian [66-61.1] myr representing the stem lineage of the Acetabularieae [58] here represented by *Acetabularia acetabulum*, *A. crenulata* and *A. dentata*. This calibration was modeled with a lognormal distribution (LogN mean=1.14, stdev=1.0, Offset=61.1). We ran the analysis for 150,000,000 generations, sampling every 1000 generations. Length of burn-in was determined by visual inspection of traces in TRACER v.1.6 [54]. The final tree was produced by TREEANNOTATOR using the “maximum clade creditability” option and mean node height, after burn-in of 1500 generations. The convergence to the stationary distribution was confirmed by inspection of the MCMC samples and of effective sample sizes (ESS should be > 200) using TRACER v1.6.0.

## Biogeographic analyses

We used the R package BIOGEOBEARS (https://cran.r-project.org/web/packages/BioGeoBEARS/index.html) [59,60] to estimate the ancestral ranges within *Ulva*. BIOGEOBEARS calculates maximum likelihood estimates of the ancestral states (range inheritance scenarios) at speciation events by modeling transitions between discrete states (biogeographical ranges) along phylogenetic branches as a function of time. Available models include a likelihood version of DIVA ([Dispersal - Vicariance Analysis 61]), Lagrange’s DEC model ([Dispersal-Extinction-Cladogenesis 62]) and BayArea [63]. Additionally, it implements the parameter “+J” that describes founder-event speciation, which is fundamental in oceanic settings [60,64].

We defined 10 geographical areas: (1) Central Indo-Pacific (CIP); (2) Eastern Indo-Pacific (EIP); (3) Tropical Atlantic (TAT); (4) Temperate Australasia (TAU); (5) Temperate Northern Atlantic (TNA); (6) Temperate Northern Pacific (TNP); (7) Temperate South Americ (TSA); (8) Western Indo-Pacific (WIP); (9) Tropical Eastern Pacific (TEP) and (10) Temperate South Africa (TSAf). According to the algae data base (http://www.algaebase.org/), the maximum number of areas that any species may occur was set to seven.

## Acknowledgements

MBB was supported by the Fundação Grupo Boticário de Proteção à Natureza (1051-20152), Conselho Nacional de Desenvolvimento Científico e Tecnológico (CNPq 306917/2009-2 to PAH) and Coordenação de Aperfeiçoamento de Pessoal de Nível Superior (CAPES/PNPD 02828/09-0 and CAPES/PNADB 2338000071/2010-61 to PAH), Brazilian Research Network on Global Climate Change and Fundação de Amparo à Pesquisa e Inovação do Estado de Santa Catarina (FAPESC). RLC was supported by a post-doctoral fellowship (SFRH/BPD/109685/2015) from FCT (Fundação para a Ciência e Tecnologia, Portugal). We thank the Portuguese FCT for CCMAR Strategic Plan PEst-C/MAR/LA0015/2011and UID/Multi/04326/2013 (RC).

## References

1. Wichard T, Charrier B, Mineur F, Bothwell JH, Clerck OD, et al. (2015) The green seaweed Ulva: a model system to study morphogenesis. Frontiers in Plant Science 6.

2. Wolf MA, Sciuto K, Andreoli C, Moro I (2012) Ulva (Chlorophyta, Ulvales) biodiversity in the North Adriatic Sea (Mediterranean, Italy): cryptic species and new introductions. J Phycol 48: 1510–1521.

3. Kozhenkova SI, Chernova EN, Shulkin VM (2006) Microelement composition of the green alga Ulva fenestrata from Peter the Great Bay, Sea of Japan. Rus J Mar Biol 32: 289–296.

4. Scherner F, Ventura R, Barufi JB, Horta PA (2013) Salinity critical threshold values for photosynthesis of two cosmopolitan seaweed species: providing baselines for potential shifts on seaweed assemblages. Marine environmental research 91: 14–25.

5. Flagella MM, Andreakis N, Hiraoka M, Verlaque M, Buia MC (2010) Identification of cryptic Ulva species (Chlorophyta, Ulvales) transported by ballast water. J Biol Res- Thessaloniki 13: 1–11.

6. Liu Q, Yu R-C, Yan T, Zhang Q-C, Zhou M-J (2015) Laboratory study on the life history of bloom-forming Ulva prolifera in the Yellow Sea. Estuarine, Coastal and Shelf Science 163: 82–88.

7. Hayden HS, Blomster J, Maggs CA, Silva PC, Stanhope MJ, et al. (2003) Linnaeus was right all along: Ulva and Enteromorpha are not distinct genera. Eur J Phycol 38: 277–294.

8. Gao G, Zhong Z, Zhou X, Xu J (2016) Changes in morphological plasticity of Ulva prolifera under different environmental conditions: A laboratory experiment. Harmful algae 59: 51–58.

9. Loughnane CJ, McIvor LM, Rindi F, Stengel DB, Guiry MD (2008) Morphology, rbcL phylogeny and distribution of distromatic Ulva (Ulvophyceae, Chlorophyta) in Ireland and southern Britain. Phycologia 47: 416–429.

10. Guiry MD, Guiry GM (2018) AlgaeBase. http://wwwalgaebaseorg/ World-wide electronic publication, National University of Ireland, Galway.

11. Perrot T, Rossi N, Ménesguen A, Dumas F (2014) Modelling green macroalgal blooms on the coasts of Brittany, France to enhance water quality management. Journal of Marine Systems 132: 38–53.

12. Kazi MA, Kavale MG, Singh VV (2016) Morphological and molecular characterization of Ulva chaugulii sp. nov., U. lactuca and U. ohnoi (Ulvophyceae, Chlorophyta) from India. Phycologia 55: 45–54.

13. Hayden HS, Waaland JR (2004) A molecular systematic study of Ulva (Ulvaceae, Ulvales) from the northeast Pacific. Phycologia 43: 364–382.

14. Hofmann LC, Nettleton JC, Neefus CD, Mathieson AC (2010) Cryptic diversity of Ulva (Ulvales, Chlorophyta) in the Great Bay Estuarine System (Atlantic USA): introduced and indigenous distromatic species. Eur J Phycol 45: 230–239.

15. Guidone M, Thornber C, Wysor B, O’Kelly CJ (2013) Molecular and morphological diversity of Narragansett Bay (RI, USA) Ulva (Ulvales, Chlorophyta) populations. J Phycol 49: 979–995.

16. Heesch S, Broom JE, Neill KF, Farr TJ, Dalen JL, et al. (2009) Ulva, Umbraulva and Gemina: genetic survey of New Zealand taxa reveals diversity and introduced species. Eur J Phycol 44: 143–154.

17. Kraft LGK, Kraft GT, Waller RF (2010) Investigations into southern Australian Ulva (Ulvophyceae; Chlorophyta) taxonomy and molecular phylogeny indicate both cosmopolitanism and endemic cryptic species. J Phycol 46: 1257–1277.

18. O’Kelly CJ, Kurihara A, Shipley TC, Sherwood AR (2010) Molecular assessment of ulva spp. (Ulvophyceae, Chlorophyta) in the Hawaiian Islands. J Phycol 46: 728–735.

19. Spalding HL, Conklin KY, Smith CM, O’Kelly CJ, Sherwood AR, et al. (2016) New Ulvaceae (Ulvophyceae, Chlorophyta) from mesophotic ecosystems across the Hawaiian Archipelago. J Phycol 52: 40–53.

20. Shimada S, Yokoyama N, Arai S, Hiraoka M (2008) Phylogeography of the genus Ulva (Ulvophyceae, Chlorophyta), with special reference to the Japanese freshwater and brackish taxa. J App Phycol 20: 979–989.

21. Azevedo CAAd, Cassano V, Júnior PAH, Batista MB, de Oliveira MC (2015) Detecting the non-native Grateloupia turuturu (Halymeniales, Rhodophyta) in southern Brazil. Phycologia 54: 451–454.

22. Sissini MN, de Barros Barreto MBB, Széchy MTM, de Lucena MB, Oliveira MC, et al. (2016) The floating Sargassum (Phaeophyceae) of the South Atlantic Ocean–likely scenarios. Phycologia 56: 321–328.

23. Bernardes Batista M, Batista Anderson A, Franzan Sanches P, Simionatto Polito P, Lima Silveira TC, et al. (2018) Kelps’ Long-Distance Dispersal: Role of Ecological/Oceanographic Processes and Implications to Marine Forest Conservation. Diversity 10: 11.

24. Graham MH, Kinlan BP, Grosberg RK (2009) Post-glacial redistribution and shifts in productivity of giant kelp forests. Proc R Soc Lond B Biol Sci: rspb20091664.

25. hu ZM, Uwai S, yu SH, Komatsu T, Ajisaka T, et al. (2011) Phylogeographic heterogeneity of the brown macroalga Sargassum horneri (Fucaceae) in the northwestern Pacific in relation to late Pleistocene glaciation and tectonic configurations. Mol Ecol 20: 3894–3909.

26. Gelman A, Rubin D (1992) A single series from the Gibbs sampler provides a false sense of security. In: Bernardo JM, Berger J, Dawid AP, Smith AFM, editors. Bayesian Statistics New York: Oxford University Press. pp. 625–631.

27. Verbruggen H, Ashworth M, LoDuca ST, Vlaeminck C, Cocquyt E, et al. (2009) A multi- locus time-calibrated phylogeny of the siphonous green algae. Mol Phylogenetics Evol 50: 642–653.

28. Ree RH, Sanmartin I (2018) Conceptual and statistical problems with the DEC+J model of founder-event speciation and its comparison with DEC via model selection. J Boigeogr 45: 741–749.

29. Lewin RA (1981) Three species concepts. Taxon: 609–613.

30. Tan IH, Blomster J, Hansen G, Leskinen E, Maggs CA, et al. (1999) Molecular phylogenetic evidence for a reversible morphogenetic switch controlling the gross morphology of two common genera of green seaweeds, Ulva and Enteromorpha. Mol Biol Evol 16: 1011–1018.

31. Wan AHL, Wilkes RJ, Heesch S, Bermejo R, Johnson MP, et al. (2017) Assessment and Characterisation of Ireland’s Green Tides (Ulva Species). PLoS ONE 12: e0169049.

32. Hiraoka M, Ichihara K, Zhu W, Shimada S, Oka N, et al. (2017) Examination of species delimitation of ambiguous DNA-based Ulva (Ulvophyceae, Chlorophyta) clades by culturing and hybridisation. Phycologia 56: 517–532.

33. Kirkendale L, Saunders GW, Winberg P (2013) A molecular survey of Ulva (Chlorophyta) in temperate Australia reveals enhanced levels of cosmopolitanism. J Phycol 49: 69–81.

34. Kerswell AP (2006) Global biodiversity patterns of benthic marine algae. Ecology 87: 2479–2488.

35. Keith SA, Kerswell AP, Connolly SR (2014) Global diversity of marine macroalgae: environmental conditions explain less variation in the tropics. Global ecology and biogeography 23: 517–529.

36. Briggs JC (2003) Marine centres of origin as evolutionary engines. J Biogeogr 30: 1–18.

37. Norton T (1992) Dispersal by macroalgae. British Phycological Journal 27: 293–301.

38. CABI. Invasive Species Compendium. Wallingford UCIwcoi (2018) CABI - Invasive Species Compendium. CAB International. http://www.cabi.org/isc. Wallingford, UK.

39. Flagella MM, Verlaque M, Soria A, Buia MC (2007) Macroalgal survival in ballast water tanks. Mar Pollut Bull 54: 1395–1401.

40. Joly AB (1965) Flora marinha do litoral norte do estado de São Paulo e regiões circunvizinhas. Bol Fac Filos Cienc Let 21: 1–267.

41. Manhart JR (1994) Phylogenetic analysis of green plant rbcL sequences. jMolecular phylogenetics and evolution 3: 114-127 %@ 1055–7903.

42. Lis JT, Schleif R (1975) Size fractionation of double-stranded DNA by precipitation with polyethylene glycol. Nucleic acids research 2: 383-390 %@ 1362–4962.

43. Puillandre N, Lambert A, Brouillet S, Achaz G (2012) ABGD, Automatic Barcode Gap Discovery for primary species delimitation. Mol Ecol 21: 1864–1877.

44. Zhang J, Kapli P, Pavlidis P, Stamatakis A (2013) A general species delimitation method with applications to phylogenetic placements. Bioinformatics 29: 2869–2287.

45. Kapli P, Lutteropp S, Zhang J (2017) Multi-rate Poisson tree processes for single-locus species delimitation under maximum likelihood and Markov chain Monte Carlo. Bioinformatics 33: 1630–1638.

46. Pons J, Barraclough T, Gomez-Zurita J, Cardoso A, Duran D, et al. (2006) Sequence-based species delimitation for the DNA taxonomy of undescribed insects. Syst Biol 55: 595–609.

47. R Core Team (2018) R: a language and environment for statistical computing. R Foundation for Statistical Computing. In: Computing RFfS, editor. 3.5 ed. Vienna, Austria.

48. Bouckaert R, Heled J, Kühnert D, Vaughan T, Wu C-H, et al. (2014) BEAST 2: a software platform for Bayesian evolutionary analysis. PLoS Comp Biol 10: e1003537.

49. Katoh K, Toh H (2010) Parallelization of the MAFFT multiple sequence alignment program. Bioinformatics 26: 1899–1900.

50. Akaike H (1973) Information theory as an extension of the maximum likelihood principle. In: Csaksi BNPaF, editor. 2nd International Symposium on Information Theory. Budapest, Hungary: Akademiai Kiado. Pp. 267–281

51. Schliep KP (2011) Phangorn: phylogenetic analysis in R. Bioinformatics 27: 592–593.

52. Guindon S, Gascuel O (2003) A simple, fast and accurate algorithm to estimate large phylogenies by maximum likelihood. Syst Biol 52: 696–704.

53. Ronquist F, Teslenko M, van der Mark P, Ayres DL, Darling A, et al. (2012) MrBayes 3.2: efficient Bayesian phylogenetic inference and model choice across a large model space. Syst Biol 61: 539–542.

54. Rambaut A, Suchard MA, Xie D, Drummond AJ (2014) Tracer version 1.6, Available from http://beast.bio.ed.ac.uk/Tracer.

55. Drummond AJ, Suchard MA, Xie D, Rambaut A (2012) Bayesian phylogenetics with BEAUti and the BEAST 1. Mol Biol Evol 29: 1969–1973.

56. Kenrick P, Li C-S (1998) An early, non-calcified, dasycladalean alga from the Lower Devonian of Yunnan Province, China. Rev Palaeobot Palynol 100: 73–88.

57. Cocquyt E (2009) Phylogeny and molecular evolution of green algae: Ghent University. 167 pp. p.

58. Morellet L, Morellet J (1922) Nouvelle contribution à l’étude des Dasycladacées tertiaires.- Paris. 35 p. p.

59. Matzke NJ (2013) BioGeoBEARS: BioGeography with Bayesian (and Likelihood) Evolutionary Analysis in R Scripts. R package, version 0.2.1, published July 27,2013 at: http://cran.r-project.org/package=BioGeoBEARS.

60. Matzke NJ (2013) Probabilistic historical biogeography: new models for founder-event speciation, imperfect detection, and fossils allow improved accuracy and model-testing. Front Biogeogr 5.

61. Ronquist F (1997) Dispersal-vicariance analysis: A new approach to the quantification of historical biogeography. Syst Biol 46: 195–203.

62. Ree RH, Smith SA (2008) Maximum likelihood inference of geographic range evolution by dispersal, local extinction, and cladogenesis. Syst Biol 57: 4–14.

63. Landis MJ, Matzke NJ, Moore BR, Huelsenbeck JP (2013) Bayesian Analysis of Biogeography when the Number of Areas is Large. Syst Biol 62: 789–804.

64. Matzke NJ (2014) Model selection in historical biogeography reveals that founder-event speciation is a crucial process in island clades. Syst Biol 63: 951–970.

65. Spalding MD, Fox HE, Allen GR, Davidson N, Ferdaña ZA, et al. (2007) Marine ecoregions of the world: a bioregionalization of coastal and shelf areas. AIBS Bulletin 57: 573–583.

